# Glucose availability impacts proteotoxic stress in *Caenorhabditis elegans*

**DOI:** 10.1101/763060

**Authors:** Landon Gatrell, Whitney Wilkins, Priya Rana, Mindy Farris

## Abstract

Alterations in protein folding may lead to aggregation of misfolded proteins, which is strongly correlated with neurotoxicity and cell death. Protein aggregation has been shown as a normal consequence of aging, but it is largely associated with age-related disease, particularly neurodegenerative diseases like Huntington disease (HD). Huntington disease is caused by a CAG repeat expansion in the *huntingtin* gene and serves as a useful model for neurodegeneration due to its strictly genetic origin. Research in the model organism *Caenorhabditis elegans* suggests that glucose protects against cell stress, including proteotoxicity related to aggregation, despite the well-known, lifespan-shortening effects of glucose. We hypothesized that glucose could be beneficial by alleviating energy deficiency, a well-characterized phenomenon in HD, or by upregulating stress resistance pathways. We used *C. elegans* expressing polyglutamine repeats to quantify lifespan, motility, reproduction, learning, and activity of succinate dehydrogenase (SDH), with and without glucose, to identify the role of glucose in proteotoxicity and neuroprotection. Our data show HD worms on glucose plates exhibited shorter lifespans, no change in motility, learning, or SDH product formation, but had altered reproductive phenotypes similar to dietary restriction. Additionally, worms expressing toxic polyglutamine repeats were unable to learn association of food with a neutral odorant. We also observed tissue-specific differences; polyglutamine appeared to be slightly more toxic to muscle cells than neurons. Rather than increasing energy production, glucose appeared to decrease mitochondrial metabolism, as SDH formation decreases with added glucose. Future work investigating glucose-mediated neuroprotection should focus on connecting metabolism, sirtuin activation, and DAF-16 activation.

## Introduction

Protein aggregates are a shared phenotype of many fatal neurodegenerative diseases including polyglutamine disorders such as Huntington disease (HD) [1]. The exact nature of cytotoxicity and neuronal death in HD and other neurodegenerative diseases is currently unknown, but many hypotheses have been proposed, including: loss-of-function and/or gain-of-function mutations, excitotoxicity, oxidative stress, aberrant gene expression, and apoptosis [2–5]. HD provides a convenient model for neurodegeneration research, as it is strictly a genetic disease caused by a specific mutation: a CAG trinucleotide repeat expansion in the gene encoding the huntingtin protein (HTT) [6]. HTT is expressed ubiquitously, though most abundantly in neurons, and is found within the nucleus and cytoplasm of the cell. Similarly, HD-afflicted neurons exhibit intranuclear and cytosolic aggregates containing mutant HTT, ubiquitin, and other proteins [7]. What role these aggregates play in neurotoxicity is unclear, with competing hypotheses of deleterious and even protective functions [8, 9]. Perhaps the most interesting aspect of neurodegenerative disease is the delayed onset of symptoms until later in life, despite lifelong and ubiquitous expression of the mutant protein [10]. This has led to research linking mechanisms of aging to aggregation diseases.

Loss of proteostasis is an inherent part of aging in wild-type systems and age-related protein aggregation has been observed in the wild-type strain of the model organism *Caenorhabditis elegans* [11, 12]. Down-regulation of proteostasis is of particular importance to neurodegenerative research and is currently hypothesized to be a contributing factor to the age-related progression of neurodegeneration [13]. Another major feature of aging is an individual’s interaction with the environment; specifically, diet has been shown to be a major modulator of aging. A common feature of aging research deals with dietary restriction (DR) and glucose enrichment (GE), which lengthen and shorten lifespan respectively. In *C. elegans*, the transcription factors DAF-16 (dauer abnormal factor 16) and HSF-1 (heat shock factor 1) play critical roles in regulating and effecting these phenotypes and both are affected by the activity of the insulin/IGF signaling pathway [14]. Upregulating the insulin/IGF pathway via GE results in deactivation of DAF-16 and HSF-1, decreased lifespans, and accelerated aging [14]. Genetic knockdown of DAF-16 and HSF-1 also accelerates aging and animals display aging phenotypes similar to GE [14]. Conversely, the lifespan extension of some DR techniques downregulates the insulin/IGF pathway and requires DAF-16 and HSF-1 [14].

Despite shortening lifespan, GE has recently been shown to increase resistance to proteotoxicity in *C. elegans*, as reduced neurodegeneration was seen in worms expressing long polyglutamine tracts (128Q) or TDP-43 when fed high glucose diets [15]. The insulin/IGF pathway appears to be involved in this, as DAF-16 and HSF-1 were required for glucose-mediated neuroprotection [15]. Glucose feeding theoretically increases energy availability, but it is unclear whether this actually leads to increased energy production. Glucose addition has been shown to cause initial increase in mitochondrial function, followed by decrease, while restriction of glucose has been shown to promote mitochondrial metabolism [16, 17]. It is possible that cells with high glucose diets downregulate energy production pathways to prevent increased ROS accumulation. HTT seems to have an indirect role in energy production in neurons, and HD neurons display many symptoms of energy deficiency (reviewed in [3]). For example, cells expressing a polyglutamine fragment have reduced expression and activity of succinate dehydrogenase (SDH)/Complex II in the electron transport chain [18].

To gain a better understanding of the phenotypic effects of glucose-mediated neuroprotection, we asked if glucose enrichment affected phenotypes associated with HD, such as motility and lifespan, in shorter polyglutamine models of *C. elegans*. Although *C. elegans* do not contain an ortholog to human HTT, expression of polyglutamine repeats alone has been demonstrated to cause aggregate formation and toxic effects resembling polyglutamine disease phenotypes [19]. We measured lifespan, motility, reproduction, SDH product formation, and learning in model organisms expressing expanded (40Q), non-toxic (24Q), and threshold (35Q) polyglutamine repeats. We found that while the presence of glucose failed to improve the resistance to proteotoxic stressors for poly-Q worms, through the simultaneous use of multiple different polyglutamine strains of *C. elegans*, some tissue-specific differences were shown. Glucose effects on reproduction were surprising, as we saw no significant effect of glucose on the brood size or egg-laying pattern of N2 animals, contrary to that seen in studies using lower concentrations of glucose [14, 15]. Glucose effects on two polyglutamine strains, however, mirror the reduced brood size and delayed egg-laying phenotype of worms under DR conditions [20, 21]. Using a butanone food-association assay, we showed HD worms to be deficient in learning, even as 1-day-old adults, before other proteotoxic phenotypes are apparent. Glucose failed to have a measurable effect on this learning phenotype or on SDH activity.

## Materials and Methods

### *C. elegans* Strains

*Caenorhabditis elegans* strains were acquired from the *Caenorhabditis* Genetics Center at the University of Minnesota. *C. elegans* strains used in this project were: the N2 Bristol (wild-type), AM101 (*rmls110*), AM138 (*rmls130*), AM140 (*rmls132*), and AM141 (*rmls133*) strains. The polyglutamine strains (AMXXX) express a construct that contains a polyglutamine repeat of varying length tagged by yellow fluorescent protein (YFP), further differentiated by the promotor-driven tissue localization. The AM101 strain has a length of 40Q and the construct is expressed pan-neuronally. The AM138 strain has a length of 24Q, expressed in the body-wall muscle cells. The AM140 strain has a length of 35Q and is expressed in the body-wall muscle cells; interestingly, the AM140 strain’s polyglutamine fragment is on the verge of the toxic threshold and only older worms show aggregation. The AM141 strain has a length of 40Q and is expressed in the body-wall muscle cells.

Strains were maintained on Nematode Growth Medium (NGM) plates spotted with 250-500μL of the OP50 strain of *E. coli* at 20°C. Prior to experiments, worms were checked under a fluorescent microscope for YFP expression. For experiments, worms were moved onto new NGM plates or NGM plates with 250mM glucose added (referred to as glucose plates, G) as appropriate. When not being manipulated for experiments, worms were kept at 20°C. Bacterially-starved worms or contaminated plates were not used for experiments.

### Lifespan Assay

The lifespans of the *C. elegans* strains were measured with (G) or without (NGM) 250mM glucose added. Polyglutamine (polyQ) and N2 worms at the larval L4 stage were transferred from general populations to new, clean NGM plates with or without 250mM glucose (G) spotted with 250μL of the OP50 strain of *E. coli*. Surviving worms were scored every other day until all experimental worms were dead. Surviving, reproductively active worms were moved to new plates of the same condition to exclude progeny from analyses. Worms were considered dead when they no longer responded to gentle tactile stimulation. Deaths were separated into two major categories: natural and unnatural. Unnatural deaths were accounted for in the survival analysis software.

Kaplan-Meier survival curves were generated using JMP software version 12. The log-rank test was used to test for an overall difference between groups at a 0.05 alpha level. The log rank test was also used for pairwise comparisons of interest with a Bonferroni adjusted alpha. There were four comparisons of interest (N2/NGM vs polyQ/NGM; N2/NGM vs N2/glucose; polyQ/NGM vs polyQ/glucose; N2/glucose vs polyQ/glucose) which yielded an adjusted alpha of 0.0125 for pairwise comparisons (0.05/4 = 0.0125).

### Motility Assay

The movement of N2 and polyQ worms was measured on plates with (G) or without (NGM) glucose added. Worms at the L4 larval stage were transferred onto new NGM or glucose plates as appropriate. Starting on day 1 of adulthood, the motility of the worms was measured every other day until day 13, at which point worms become predominantly sedentary. Motility was classified as the number of times the tail tip reached the crest of the wave pattern; this frequency was measured by experimenter observation via microscopy for 1 minute. Movements in reverse were also included in data collection. Worms were undisturbed on the plates prior to measurement to relate energy output (motility) to energy availability and were transferred onto new plates post-measurement to exclude progeny. This protocol therefore differed from other protocols that measure paralysis, touch sensitivity, and escape behavior because all of these require experimenter intervention by tactile stimulation.

Significance of the effects of the model was tested using a likelihood ratio test with R software version 3.5.1. Pairwise comparisons were tested using simultaneous tests for general linear hypotheses with an adjusted alpha. Data are from 3 separate trials, with n=10 worms per strain for each trial.

### Reproduction Assay

The reproduction of individual N2 and polyQ worms was measured on NGM and glucose plates over time. A single worm was placed on an NGM or glucose plate spotted with 250μL of OP50 *E. coli*. Every day thereafter, the worm was transferred onto a new plate of the same condition. The progeny on the old plates were counted after a hatching period of 24 hours. Only viable progeny were scored for this experiment; unhatched eggs were excluded. This continued until no progeny were found on the plate for 2 consecutive days. Data are from 3 separate trials, with n=10 worms per strain for each trial.

### SDH Product Formation Assay

SDH product formation was measured *in vitro* using a modified version of published methods [18]. To extract whole protein, L4 worms matured and reproduced on NGM or glucose plates for 3-4 days, generating a semi-synchronized population. The progeny matured until L4s or young adults were present in high numbers. Worms were washed off the plates using chilled SDH extraction buffer (20mM Tris-HCl, pH 7.2; 250mM saccharose; 2mM EGTA; 40mM KCl; 1mg/mL BSA). The worms were gently centrifuged at 82 x g for 1 minute and washed to remove bacteria. Protease inhibitor (5μL of Thermo-Fisher Halt™ Protease Inhibitor Single Use Cocktail; 1mM AEBSF, 800nM aprotinin, 50μM bestatin, 20μM leupeptin, 10μ pepstatin) was added, and then worms were ground on ice for 5 minutes using a tephlon grinder and an overhead stirrer. The lysate was centrifuged at 735 x g for 5 minutes to remove cell debris. The concentration of whole protein was measured using the Bradford assay using BSA as a standard.

To measure the activity of SDH in the lysate, SDH activity buffer (100mM Tris-HCl, pH 8.3; 1mM EDTA; 20mM succinate) was added in equal volume to the lysate. A final concentration of 2mM iodonitrotetrazolium chloride was added to start the reaction.

Iodonitrotetrazolium (INT) is an electron acceptor and produces a red dye upon reduction. The reaction progressed for approximately 60 minutes at 35°C and dye formation was measured using a Nanodrop spectrophotometer at 490nm, with three replicates measured per reaction.

SDH product formation was calculated by standardizing the absorbance of INT by the concentration of the lysate. Product formation was analyzed with JMP software version 12 using a two-way ANOVA at a 0.05 alpha level, with Tukey’s HSD test for *post-hoc* analysis.

### Learning Assay

The ability of N2 and polyQ worms to associate a normally neutral odorant (butanone) to food (*E. coli*) was measured after exposure to the odorant in the presence of food. Parent worms were transferred to new NGM or glucose plates spotted with 500μL of OP50 *E. coli* and allowed to reproduce for 8-16 hours, then removed so that progeny were age synchronized. The learning assay was performed according to previously described methods [22].

Chemotaxis index was scored by the number of worms at each location and calculated as: CI = (butanone – EtOH)/(total – origin). Statistical significance was analyzed with JMP software version 12 using a two-way ANOVA.

## Results

### Lifespan

To determine if glucose is indeed protective against proteotoxicity, the lifespan of wild-type and polyglutamine (polyQ) strains of *C. elegans* were measured under NGM and glucose conditions. If glucose is protective, then the addition of glucose should increase the lifespan of worms expressing toxic polyglutamine repeats (40Q). Since glucose is known to shorten the lifespan of wild-type worms, the lifespans of N2 and of worms expressing non-toxic or threshold polyglutamine repeats (24Q and 35Q) should be shortened compared to their respective NGM lifespans.

As expected, the presence of a toxic polyglutamine repeat did shorten the lifespan of *C. elegans*. Although the lifespan of the pan-neuronal 40Q worms on NGM was not significantly shortened compared to N2 (Figures 1A,B; log-rank test, p = 0.0137), there was a strong trend toward this. The lifespan of the body-wall muscle 40Q strain on NGM plates was significantly shortened (Figure 1C; log-rank test, p < 0.0001). The 24Q worms had an increased lifespan on NGM plates compared to N2/wildtype (Figure 1D; log-rank test, p = 0.003), while the 35Q worms exhibited a reduced lifespan on NGM compared to the N2 lifespan (Figure 1E; log-rank test, p < 0.0001).

**Fig 1.**
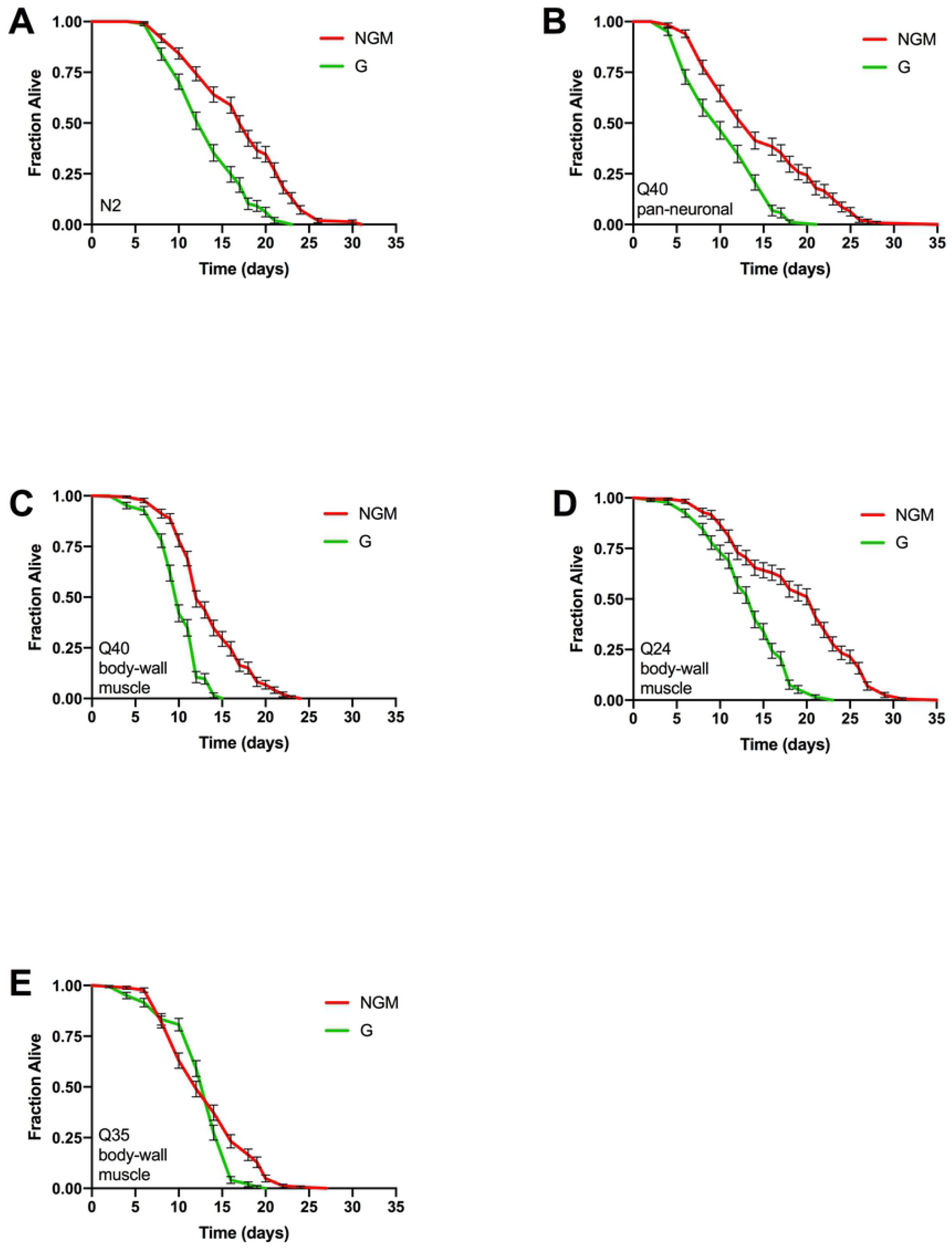
Glucose did not extend lifespans of transgenic worms expressing polyglutamine constructs. **(A)** N2 lifespan without (NGM) or with (G) glucose. **(B,C,D,E)** Lifespans of pan-neuronal 40Q, body-wall muscle 40Q, body-wall muscle 24Q, and body-wall muscle 35Q, respectively, without and with glucose. Averages are shown from three independent experiments for each strain & condition. Error bars indicate SEM.

Exposure to 250mM glucose shortened the average lifespan of *C. elegans* by approximately 3-5 days in the N2, 40Q, and 24Q strains (Figures 1A-D). The 35Q strain was the only exception to this; the glucose lifespan was not significantly different than the NGM lifespan (Figure 1E; p = 0.024).

Unexpectedly, the lifespan of toxic polyQ-afflicted worms with glucose was reduced compared to N2 worms with glucose. The lifespan of pan-neuronal 40Q worms with glucose was reduced compared to both the same strain on NGM and N2 on glucose (log-rank test, p < 0.0001 for both comparisons). The body-wall muscle 40Q on glucose was shorter than the same strain on NGM and N2 on glucose (log-rank test, p < 0.0001 for both comparisons). While the 24Q lifespan was shortened with glucose enrichment compared to on NGM, this was not significantly different from the N2/glucose lifespan (log-rank test, p < 0.0001 and p = 0.858 respectively). Interestingly, the 35Q strain was the only strain that did not exhibit a shorter lifespan on glucose plates (log-rank test, p = 0.0249). This lifespan was also not significantly different from the N2/glucose lifespan (log-rank test, p = 0.9129), similar to the 24Q strain. No lifespan-extending, protective effect of glucose was observed in any of the four polyglutamine strains, contrary to expected results.

Overall, polyglutamine shortens the lifespan of *C. elegans*; however, we observed that repeats below the toxic threshold (24Q) increase the lifespan and repeats at the toxic threshold (35Q) decrease lifespan. The addition of glucose shortened the lifespans of the 40Q strains, contrary to expectations. Glucose shortened the lifespan of the N2 and 24Q strains, but not the 35Q strain. The lifespan shortening effect of glucose in the N2 and 24Q strains and the lifespan of the 35Q strain on glucose were not different from each other.

### Motility

The spontaneous movement of wild-type and polyglutamine strains were measured to test the hypothesis that glucose affects polyQ phenotypes based on increased energy availability. Toxic polyQ worms should be less motile than wild-type due to energy deficiency, and glucose enrichment could increase the energy available and provide protection. This assay was designed to observe energy-output and relate output to energy fluctuations; therefore, worms were left undisturbed on the plates, contrary to measurements of the ability to move. This protocol assumed that the amount of spontaneous movement (without experimenter intervention) reflects the abundance of energy available to spend on movement.

Spontaneous movement was significantly affected by strain (Figure 2; χ^2^ = 103.3, df = 4, p < 0.0001). The N2 strain was more motile than the 40Q, 24Q, and 35Q body-wall muscle strains (Figures 2A,C-E; p < 0.001, p = 0.007, p < 0.0001, respectively). The pan-neuronal 40Q strain was more motile than all other strains (Figure 2B; p < 0.0001 across all comparisons). The motility of 40Q, 24Q, and 35Q body-wall muscle worms was not significantly different across strains.

**Fig 2.**
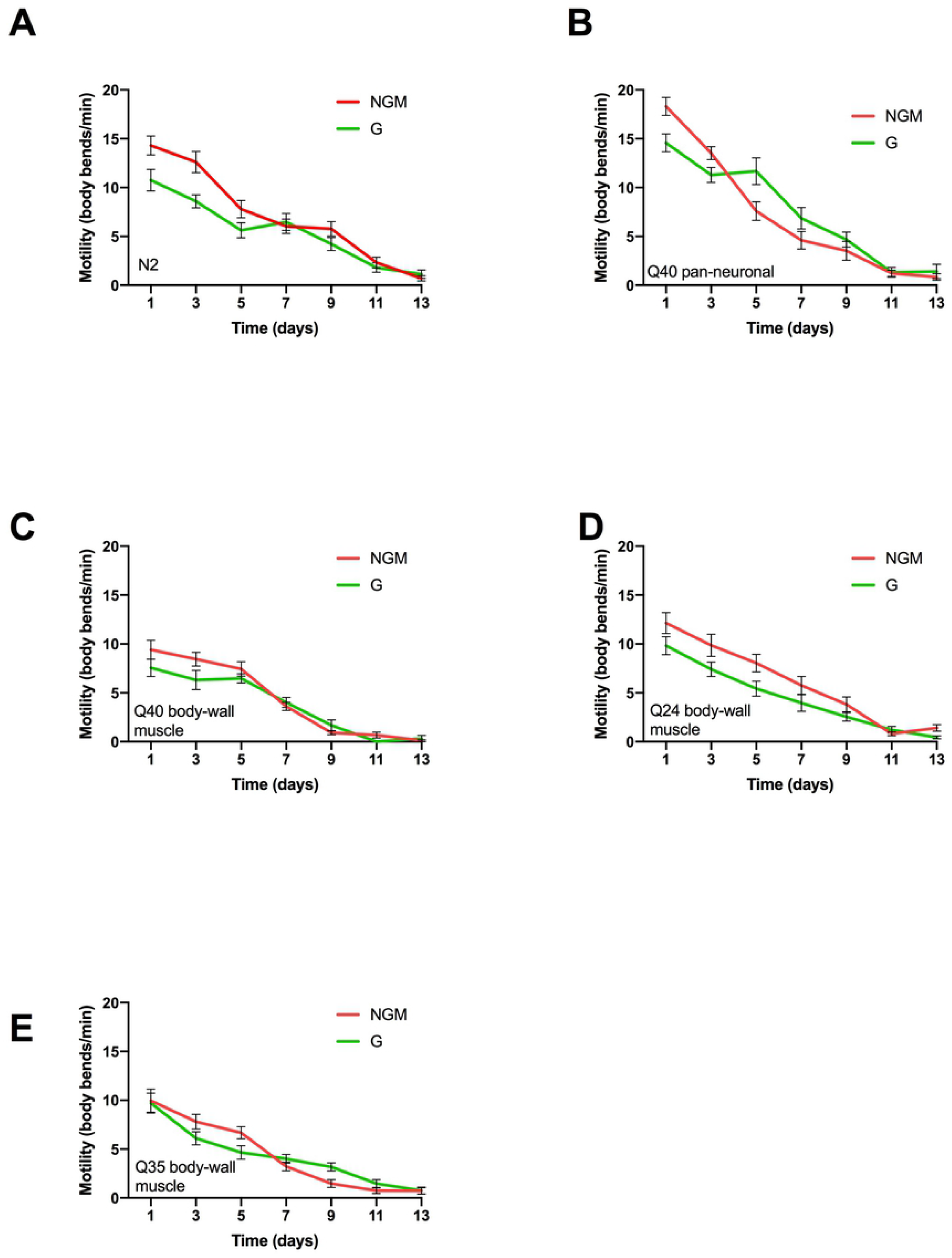
Glucose did not affect motility of transgenic worms expressing polyglutamine constructs. **(A)** N2 motility without (NGM) or with (G) glucose. **(B,C,D,E)** Motility of pan-neuronal 40Q, body-wall muscle 40Q, body-wall muscle 24Q, and body-wall muscle 35Q, respectively, without and with glucose. Averages are shown from three independent experiments for each strain & condition. Error bars indicate SEM.

Spontaneous movement was not significantly impacted by glucose (χ^2^ = 1.05, df = 1, p = 0.305), but tended to decrease motility early in adulthood, especially in the N2 strain (Figure 2).

### Reproduction

To further test how glucose affects high energy functions, the reproduction patterns of the wild-type and toxic polyQ (40Q) strains were measured. If polyglutamine proteotoxicity results in energy deficiency and this deficiency disrupts high energy functions, then polyglutamine worms should show decreased fecundity compared to N2. Furthermore, if neuroprotection occurs through increased availability of energy, then the presence of glucose should rescue reproductive deficiency in the 40Q strains.

The wild-type strain of *C. elegans* lays many eggs over a relatively short period of time (~300 over 7 days, Figure 3A). The addition of a toxic polyglutamine repeat, regardless of localization, decreases the brood size (Figures 3B,C). Contrary to expectations, the addition of glucose decreased the brood size in both 40Q strains (Figure 3). However, glucose also increased the reproductive lifespan of the worms, as worms on glucose plates continued to lay eggs for 1-2 days longer than worms on NGM plates, similar to effects of dietary restriction [20, 21].

**Fig 3.**
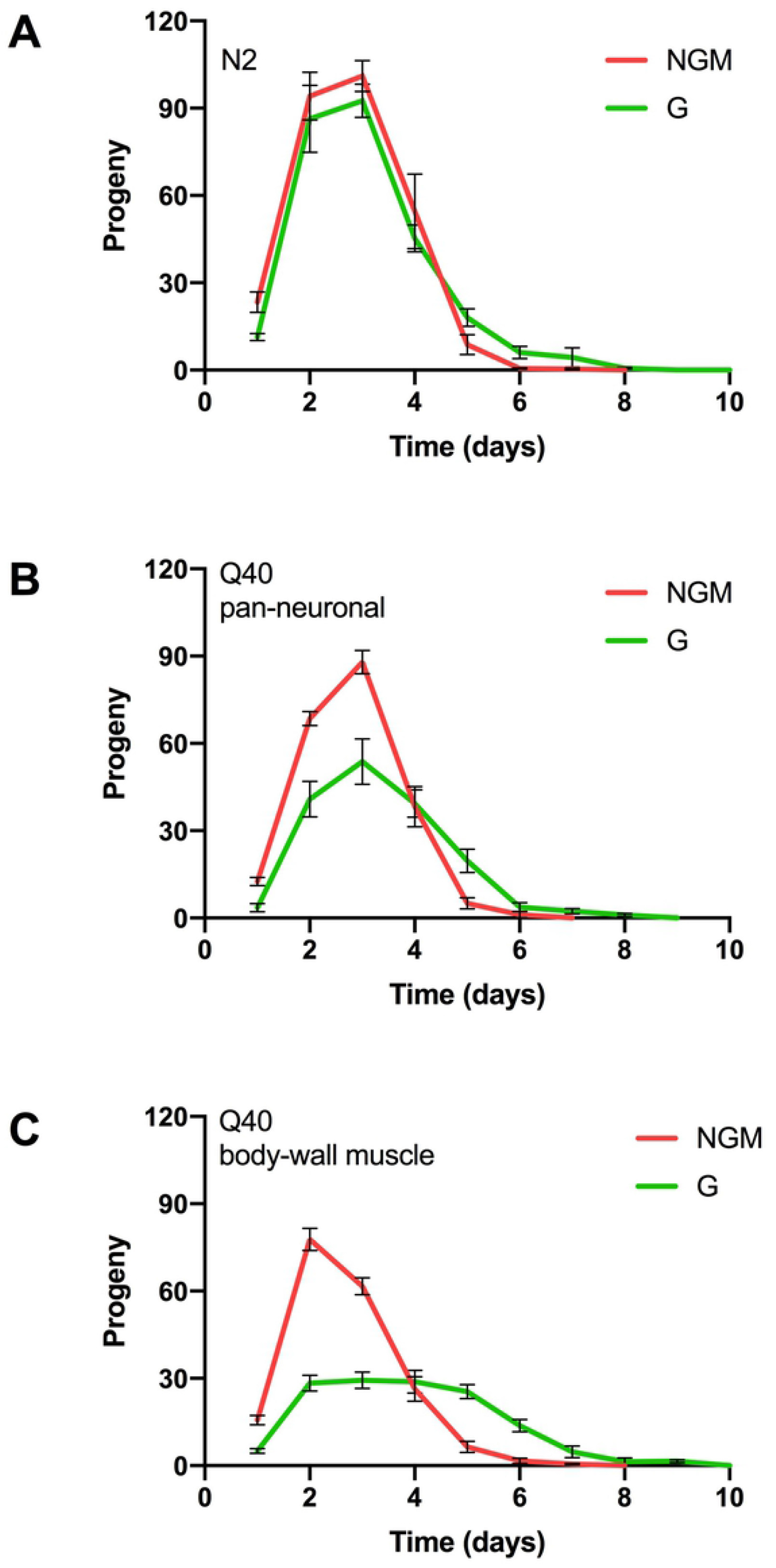
Glucose altered the egg-laying phenotype of transgenic worms expressing toxic polyglutamine constructs. **(A)** N2 reproduction without (NGM) or with (G) glucose. **(B,C)** Reproduction of pan-neuronal 40Q and body-wall muscle 40Q, respectively, without and with glucose. Averages are shown from three independent experiments for each strain & condition. Error bars indicate SEM.

### SDH Product Formation

SDH product formation was measured in young adult worms to observe how glucose affects energy deficiency in HD. We predicted the 40Q strains would show decreased SDH product formation compared to N2 due to the presence of toxic polyQ repeats. The addition of glucose was expected to increase SDH product formation based on the increased availability of energy in the N2, 24Q, and 35Q strains. If SDH deficiency in HD is the result of direct action of polyglutamine on SDH, then SDH deficiency in 40Q worms should persist despite glucose. However, if SDH deficiency is the result of energy depletion and lack of substrate, then increased glucose intake should increase SDH product formation.

SDH product formation seemed to vary widely between strains and conditions, though there was a detectable overall significant difference between groups (Figure 4; two-way ANOVA, F_9,20_ = 2.79, p = 0.026). The strain effect was the only significant effect in the model (F_4_ = 3.53, p = 0.024), although the effect of glucose verged on significance (F_1_ = 3.55, p = 0.074) with no interaction effect (F_4_ = 1.86, p = 0.15). Between strains, the product formation in 35Q worms was higher than in 40Q worms expressing the polyQ in the same tissue (p = 0.016). This was the only statistically significant pairwise comparison in the model.

**Fig 4.**
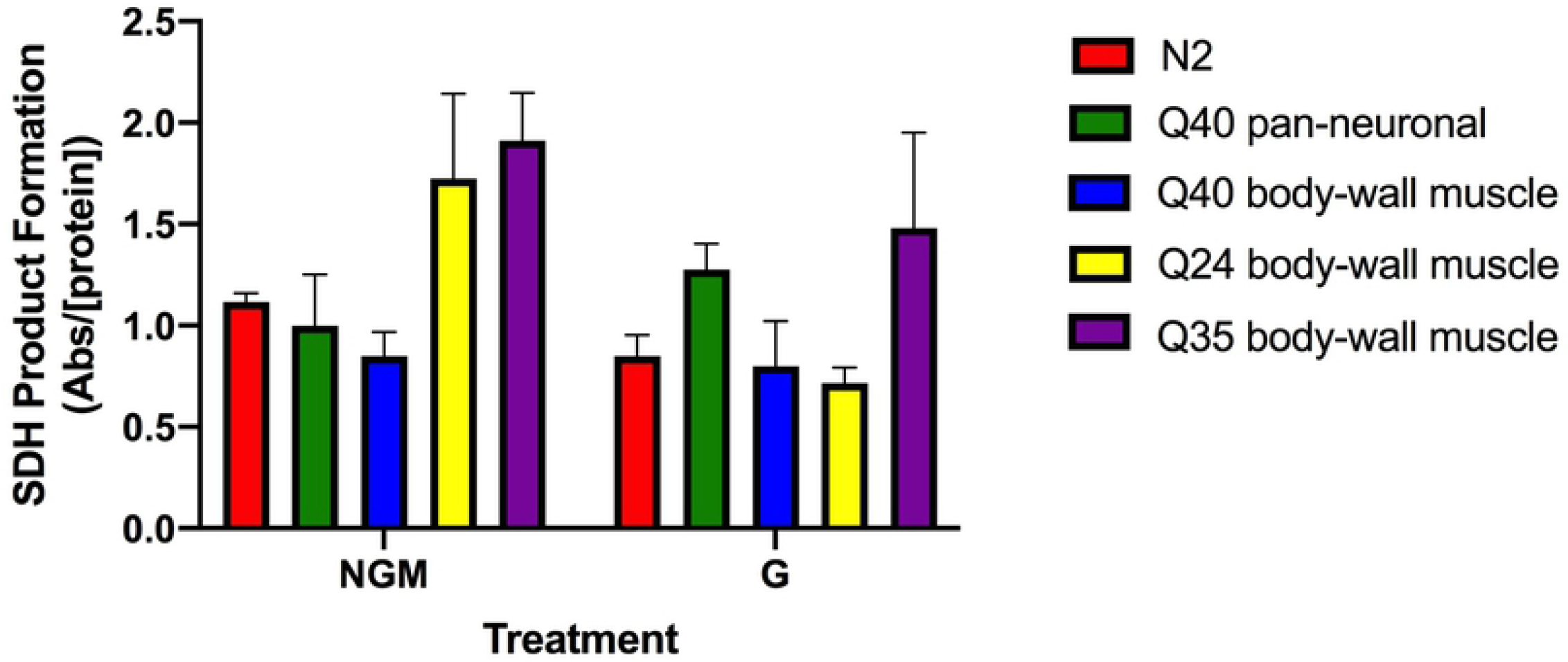
SDH product formation only changes upon addition of glucose for the non-toxic polyglutamine construct. Averages are shown from three independent experiments for each strain & condition. Error bars indicate SEM.

Although not statistically significant, some interesting trends can be seen in the data. The SDH product formation of pan-neuronal 40Q on NGM was not different from N2 on NGM. The 24Q and 35Q worms both had increased activity on NGM compared to all other strains, despite both strains carrying a non-toxic repeat. Glucose tended to decrease SDH product formation in the N2 and 24Q strains, but not in the 35Q or 40Q strains. The SDH product formation of the pan-neuronal 40Q strain actually tended to increase with glucose addition, while the enzyme activity of body-wall muscle 40Q and 35Q strains remained the same.

### Learning

As shown previously, the N2 strain of *C. elegans* is able to learn an artificial association between food (*E. coli*) and a neutral odorant (butanone) (Figure 5). Interestingly, in the pan-neuronal 40Q strain, learning was completely abolished, indicating a novel early-life (1-day-old adult) polyQ phenotype (two-way ANOVA, F_1_ = 126.1, p < 0.0001). The addition of glucose had no impact on learning in either strain (F_1_ = 0.131, p = 0.72).

**Fig 5.**
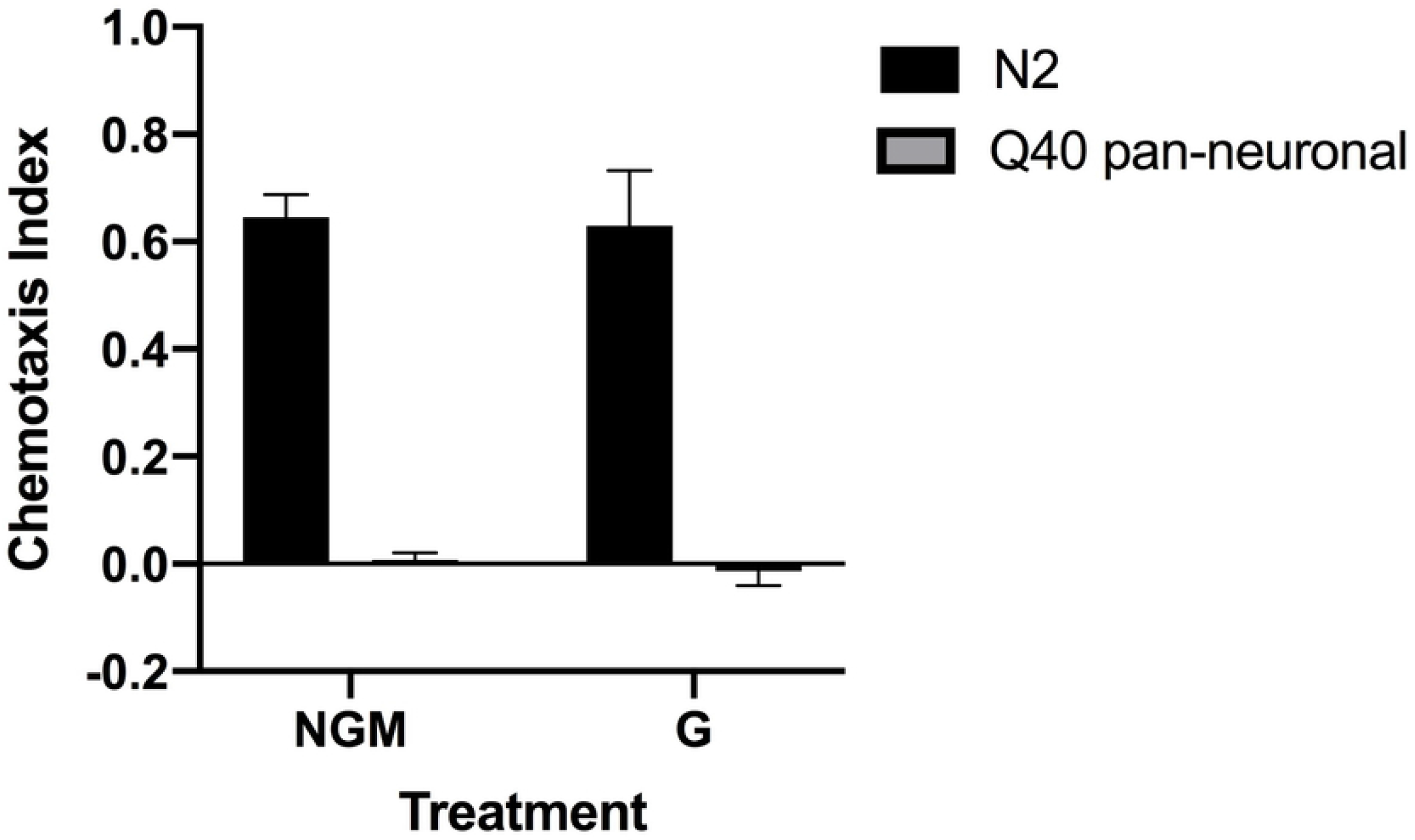
Pan-neuronal toxic polyglutamine worms show no ability to associate food with an odorant, even early in life. Glucose has no effect on the ability of these or wild-type animals to learn food association. Averages are shown from three independent experiments for each strain & condition. Error bars indicate SEM.

## Discussion

The aim of this project was to assess the glucose-dependent stress resistance of non-disease-associated (24Q), threshold (35Q), and expanded (40Q) polyglutamine models of *Caenorhabditis elegans*. To test the hypothesis that glucose is protective against proteotoxic stress, four strains of polyglutamine-expressing worms were studied under normal and high glucose diets, along with the wild-type/N2 strain. Worms with toxic polyQ expansions were expected to be deficient in lifespan, motility, reproduction, SDH product formation, and learning phenotypes compared to the wild-type strain. If glucose was protective, then high glucose diets should have improved or rescued deficiency. Furthermore, if glucose was protective based upon increased energy availability, then SDH product formation should have correlated with increased lifespan, motility, etc.

The pan-neuronal 40Q strain failed to associate butanone with *E. coli* (Figure 5; p < 0.0001). One mechanism of synaptic plasticity and learning is long-term potentiation, which functions through NMDA receptors [23]. Stimulation of these receptors recruits AMPA receptors to the post-synaptic membrane, modulates the excitability of the post-synaptic neuron, and strengthens the connection [23]. Several studies using mouse-models of HD and post-mortem human brains have shown that HD presents several dysfunctions with NMDA receptor function and AMPA receptors (extensively reviewed in [24, 25]). In *C. elegans*, NMDA and AMPA receptor homologs are present, and the worm can learn via long-term potentiation [26, 27]. The fact that pan-neuronal 40Q worms performed poorly in learning tasks suggests that polyQ disrupts long-term potentiation in *C. elegans*, similar to mouse models of HD.

NMDA and AMPA receptors are glutamate-stimulated calcium channels, known to be deficient in HD, and are heavily implicated in the excitotoxicity model of neurodegeneration [28, 24]. Calcium is an important intracellular signal and functions to control mitochondrial activity, apoptotic signaling, neurotransmitter release, etc. [29, 30]. If NMDA and AMPA receptors are deficient in *C. elegans*, then aberrant glutamate signaling may lead to calcium influxes in neuronal circuits and excitotoxicity. Calcium influx, therefore, should increase energy production, SDH activity, and ROS production, although SDH is not activated directly by Ca^2+^ [30]. However, SDH product formation was not different in the pan-neuronal 40Q strain compared to the N2 strain (Figure 4). One explanation for this could be that increased ROS production negatively regulated or reduced activity of SDH before product formation was measured [31].

Increased calcium influxes may also increase acetylcholine signaling at the neuromuscular junction, leading to increased motility in the pan-neuronal 40Q strain (Figure 2; p = 0.0001). Interestingly, this effect seems to be time-dependent; increased motility in this strain appears to be restricted to early in life, although this was not tested statistically. Aberrant calcium signaling is unlikely to affect the motility of the body-wall muscle 40Q strain because expression of the polyQ is restricted to muscles. Neurons of the body-wall muscle 40Q strain should fire normally, and thus the incoming signal at the neuromuscular junction should resemble that of wild-type. Dysfunction in the body-wall muscle 40Q strain should originate in the muscle and this is supported by the motility of the body-wall muscle 40Q strain, which was decreased compared to the pan-neuronal 40Q and N2 strains (Figure 2; p < 0.0001 for both comparisons). The increase in motility in the pan-neuronal 40Q strain and the decrease in motility in the body-wall muscle 40Q strain suggest that proteotoxicity manifests differently in neurons than in muscles.

The tissue-specific effects of polyglutamine extended to other phenotypes: lifespan of the body-wall muscle 40Q strain was slightly shorter than that of the pan-neuronal 40Q strain (Figure 1), though this effect was not statistically significant. Motility in all three muscular polyglutamine strains was decreased when compared to the pan-neuronal strain (Figure 2; p < 0.0001 for all comparisons). Brood size was slightly decreased in body-wall muscle 40Q compared to pan-neuronal 40Q (Figure 3). SDH product formation was slightly decreased in polyQ muscles compared to wild-type, but not neurons (Figure 4). Considering that both the pan-neuronal and body-wall muscle 40Q strains express a repeat of the same length, these data suggest that muscles are slightly more vulnerable to polyglutamine than neurons.

Differences in vulnerability may be due to a fundamental difference in the chaperone activity in muscles and neurons, which was investigated in *C. elegans* under heat stress [32]. Heat stress causes proteins to misfold; cells resist protein misfolding via the heat shock response and upregulation of molecular chaperones [32]. Under the control of HSF-1, muscles are more resistant to heat stress, while neurons showed a stronger recovery from heat stress. In older worms, neurons show more resistance but no recovery (similar to young muscles), while older muscles lose resistance and recovery altogether [32]. What this implies for chronic proteotoxic stress is unclear, but it could provide support for increased sensitivity to polyglutamine in muscles, leading to shorter lifespans and more deficiency. Although degeneration of specific populations of neurons is generally thought to be causative of the specific phenotypes associated with the various polyglutamine diseases, some research indicates that dysfunction is not exclusive to neurons and may explain some of the motor deficits in HD (reviewed in [33–40]).

Curiously, the 24Q strain was no more motile than the 35Q and 40Q body-wall muscle strains, which are at or above the toxic threshold (Figure 2; p = 0.29, p = 0.71 respectively). This suggests that the presence of polyQ in muscle cells, no matter the degree of toxicity, reduces movement in *C. elegans*. The 35Q strain was another peculiarity in the data; this was the only strain whose lifespan was not shortened by glucose (Figure 1; p = 0.0249). Surprisingly, the 24Q strain lived significantly longer than the N2 strain (Figure 1; p = 0.003). One potential and interesting explanation for this is that the repeat may induce cellular stress that the cell is easily able to manage via activation of stress resistance pathways, thus providing a net stress resistant phenotype (hormesis). If the 24Q strain is receiving a hormetic benefit from its non-toxic polyQ repeat, then DAF-16 and/or HSF-1 could be active in this strain under NGM conditions and may be the hormetic effector(s). One result of DAF-16 activation is stimulation of metabolism, and 24Q’s increased SDH product formation (not significant) supports the hypothesis that DAF-16 is active (Figure 4; [41–43]). In mammals, the DAF-16 homolog FOXO3a has been found to have increased nuclear localization in mouse HD cell lines [44]. FOXO3 has been shown to differentially regulate transcription in HD, connected to altered sleep and stress phenotypes appearing as early HD symptoms [45]. The mechanism associated with these early, non-motor symptoms may also be connected to the failure we observed of young (1-day-old) 40Q worms in the butanone learning assay.

SDH deficiency has been well characterized in HD models, but its origin is unclear [18, 3]. One potential explanation is that the direct action of polyglutamine affects SDH via gain of function action. SDH product formation in the body-wall muscle 40Q strain was not significantly reduced compared to wild-type, nor in the pan-neuronal 40Q strain (Figure 4). This alone suggests that polyglutamine itself is not the effector of SDH deficiency. Furthermore, glucose enrichment was not indicative of increased energy production, at least in terms of SDH, so SDH deficiency is unaffected by increased substrate availability. These results suggest that SDH deficiency arises from another source altogether. Interestingly, enzymes of the TCA cycle, such as aconitase, are particularly sensitive to ROS, which is increased in HD [3]. However, the ETC is the major generator of ROS in the cell, and the results of this study did not show increased activity of SDH/complex II in toxic polyQ strains when compared to N2. Alternatively, the subunits of SDH function separately in the TCA cycle and ETC, meaning that SDH product formation may not be indicative of complex II activity [31].

Collectively, these data make clear that glucose, at the relatively high (250mM) concentrations used in this study, is not protective in the polyglutamine models we examined. The glucose concentrations used here are possibly beyond the threshold of protection, which may explain why we did not see the protection observed in other studies, which also used longer polyglutamine constructs [15]. Glucose-mediated neuroprotection was previously found to be DAF-16-dependent and DAF-16 does have some control over metabolism [46, 15]. However, the results of this study support other studies that show glucose may not increase mitochondrial energy production, as SDH product formation was not stimulated by glucose enrichment (Figure 4; [16, 17]). Glucose-mediated neuroprotection may be due to DAF-16 activity, which would lead to interesting future questions, including what is activating DAF-16, and how the downstream targets of DAF-16 interact with factors effecting polyglutamine proteotoxicity.

## Conclusions

This study assessed the glucose-dependent stress resistance of *Caenorhabditis elegans* in response to polyglutamine proteotoxicity. As neurodegeneration poses a significant health risk, identification of both neuroprotective and neurotoxic factors is essential. *C. elegans* expressing toxic polyglutamine repeats exhibited shorter lifespans and increased deficiency, as measured by lifespan, motility, reproduction, SDH product formation, and learning. Glucose (250mM) did not improve these deficits in 24Q, 35Q, and 40Q polyglutamine models, and exacerbated decrease in lifespan. This glucose concentration is likely beyond a threshold of activation of protective mechanisms, dependent on NAD^+^ levels, sirtuin activation, and DAF-16 dependent gene expression.

## Acknowledgments

The authors would like to thank the support of the University Research Council, University of Central Arkansas, AR. We acknowledge funding from The Arkansas IDeA Network of Biomedical Research Excellence (Arkansas INBRE).

## Author Contributions

LG performed all lifespan, motility, and SDH assay experiments. LG and MF performed the reproduction experiments. WW and PR performed the learning assay experiments. MF supervised the project. LG and MF wrote the manuscript with input from all authors.

